# Adaptive introgression across semipermeable species boundaries between local *Helicoverpa zea* and invasive *Helicoverpa armigera* moths

**DOI:** 10.1101/2019.12.15.877225

**Authors:** Wendy A. Valencia-Montoya, Samia Elfekih, Henry L. North, Joana I. Meier, Ian A. Warren, Wee Tek Tay, Karl H. J. Gordon, Alexandre Specht, Silvana V. Paula-Moraes, Rahul Rane, Tom K. Walsh, Chris D. Jiggins

**Affiliations:** Department of Zoology, University of Cambridge, Cambridge, United Kingdom; CSIRO Health and Biosecurity, Australian Animal Health Laboratory, Geelong, VIC, Australia; Bio21 Institute, University of Melbourne, Parkville VIC 3052, Australia; CSIRO Land and Water, Black Mountain Laboratories, Canberra, Australia; Embrapa Cerrados, Planaltina, Federal District, Brazil; West Florida Research and Education Center, University of Florida, Jay, FL, USA

## Abstract

Hybridization between invasive and native species has raised global concern given the dramatic increase in species range shifts and pest outbreaks due to climate change, development of suitable agroecosystems, and anthropogenic dispersal. Nevertheless, secondary contact between sister lineages of local and invasive species provides a natural laboratory to understand the factors that determine introgression and the maintenance or loss of species barriers. Here, we characterize the early evolutionary outcomes following secondary contact between invasive *Helicoverpa armigera* and *H. zea* in Brazil. We carried out whole-genome resequencing of *Helicoverpa* moths from Brazil in two temporal samples: during the outbreak of *H. armigera* in 2013, and more recent populations from 2017. There is evidence for a burst of hybridization and widespread introgression from local *H. zea* into invasive *H. armigera* coinciding with *H. armigera* expansion in 2013. However, in *H. armigera*, admixture proportions were reduced between 2013 and 2017, indicating a decline in hybridization rates. Recent populations also showed shorter introgressed tracks suggesting selection against admixture. In contrast to the genome-wide pattern, there was striking evidence for introgression of a single region including an insecticide-resistance allele from the invasive *H. armigera* into local *H. zea,* which increased in frequency over time but was localized within the genome. In summary, despite extensive gene-flow after secondary contact, the species boundaries are largely maintained except for the single introgressed region containing the insecticide-resistant locus. We document the worst-case scenario for an invasive species, in which there are now two pest species instead of one, and the native species has acquired resistance to pyrethroid insecticides through introgression and hybridization, with significant implications for pest management in future population expansions and introductions of novel resistance genes from new invasive *H. armigera* populations.

**Author summary:** Secondary contact occurs when related species with non-overlapping ranges are geographically reunited. Scenarios of secondary contact have increased due to anthropogenic movement of species outside of their native range, often resulting in invasive species that successfully spread and stabilised in the new environment. This is the case for *Helicoverpa armigera*, a major agricultural pest in the Old World that has recently invaded the Americas, where it reunited with its closest relative, *H. zea*. While some authors reported hybridisation, and hypothesised about the potential emergence of novel ecotypes and the exchange of pesticide-resistant genes, these outcomes have not been tested yet. We examine these outcomes by sequencing individuals from both species in Brazil, collected in 2013 after outbreaks of *H. armigera* were reported, and individuals collected during 2017. We discovered that despite hybridisation, these moths have not collapsed into a single species nor formed new ecotypes, and that the species distinctiveness is maintained through selection against most of the foreign genotypes that cross species boundaries. However, we found that hybridisation mediated the rapid acquisition of a *H. armigera* gene conferring resistance to pyrethroids by *H. zea*. The overall decline in populations of both species during the interval covered by this study means that our results are likely to reflect the consequences of hybridization events early after invasion, despite the likely ongoing introduction of *H. armigera* genetic diversity through trade across the South American continent. Our results provide a rare example of adaptive transferral of variation right after invasion and elucidate the dynamics of insecticide resistance evolution in *H. zea*.

## Introduction

Hybridization between local and introduced species is an underestimated threat to global agriculture (Mooney and Cleland 2001; Blair and Hufbauer 2010; Paini et al. 2016). Although hybridization is often maladaptive, introgressive hybridization can occasionally introduce adaptive variants, and potentially contribute to establishment and invasiveness of the newcomer species (Currat et al. 2008; Saarman and Pogson 2015; Hall 2016; Mesgaran et al. 2016). This makes invasive hybridization a major global concern impacting natural and agricultural systems, since species introductions are predicted to continue at an accelerating rate following trade, human movement and climate change (Saarman and Pogson 2015; Paini et al. 2016). Simultaneously, invasive species offer a unique opportunity to study evolutionary outcomes following secondary contact. While secondary contact zones have played a major role in the study of reproductive isolation and speciation, it remains a challenge to disentangle the contribution of ancient and recent gene-flow to the contemporary patterns of introgression (Barton and Hewitt 1985; Abbott et al. 2013). This is primarily because ancestral range reconstruction is challenging, and so it is hard to confidently estimate the extent of historical contact between species (Barton and Hewitt 1985; Abbott et al. 2013; Saarman and Pogson 2015). However, in the case of invasive species, the geographical reunion of divergent species can be observed in “real time”. This allows us to systematically test hypotheses regarding the possible outcomes of secondary contact including fusion, reinforcement, hybrid speciation, and adaptive introgression.

The recent invasion by the Old World cotton bollworm *Helicoverpa armigera* of South America (Czepak et al. 2013), where it hybridizes with its sister species *H. zea* (Tay et al. 2013; Anderson et al. 2018), represents an excellent scenario in which to study the evolutionary trajectories following secondary contact. *Helicoverpa armigera* comprises two subspecies in the Old World, *H. armigera conferta* in Oceania (Australia, New Zealand) and *H. armigera armigera* in the rest of the Old World native range and in the invasive population in South America (Anderson et al. 2016; Anderson et al. 2018) (hereafter *H. armigera* refers to *H. armigera armigera* unless otherwise specified). *Helicoverpa zea* is present throughout the temperate and tropical regions of N. and S. America. *H. armigera* and *H. zea* are estimated to have diverged around 1.5 Mya (Behere et al. 2007; Pearce et al. 2017) as a result of a transoceanic dispersion of *H. armigera* into the New World, and thus represents a classic case of allopatric speciation (Pearce et al. 2017; Anderson et al. 2018). There is no evidence of introgression between the species during this period of divergence until the recent incursions of *H. armigera* into Brazil (Pearce et al. 2017; Anderson et al. 2018), thought to have resulted from international trade (Pearce et al. 2017; Anderson et al. 2018; Mallet 2018; Tay and Gordon 2019). There is evidence for multiple initial incursions, centred on multiple geographic locations (Tay et al. 2017; Arnemann et al. 2019). Multiple and likely ongoing introductions of *H. armigera* from the Old World mean that the genetic diversity of the New World populations is very high and possibly comparable to that in the species’ ancestral range (Tay et al. 2017; Arnemann et al. 2019; Gonçalves et al. 2019).

Although the two species are very similar morphologically and their genomes share high levels of orthology, *H. armigera* has twice the genome-wide genetic diversity and a much greater (∼10x) effective population size compared to *H. zea* (Pearce et al. 2017). In addition, *H. zea* shows a narrower host-plant range and is mainly known as a pest of corn and cotton although is also found on over 100 plants belonging to 29 families (Pearce et al. 2017; Anderson et al. 2018; Mallet 2018). *H. armigera* is more devastating to agriculture and feeds on over 300 species belonging to 68 families (10–12). These differences in host range are associated with fewer gustatory receptor and detoxification genes in the *H. zea* genome, which showed no evidence for gain of any genes after the species diverged (Pearce et al. 2017).

Another substantial phenotypic difference between *H. armigera* and *H. zea* lies in their susceptibility to insecticides widely used for pest control (Anderson et al. 2018; T.K. Walsh et al. 2018). *H. armigera* shows by far the highest number of reported cases of insecticide resistance worldwide having evolved resistance against pyrethroids, organophosphates, carbamates, organochlorines, and recently to macrocyclic lactone spinosad, and several *Bacillus thuringiensis* toxins (McCaffery 1998; Xu et al. 2005; Mahon et al. 2007; Joussen et al. 2012; Tay et al. 2015; Durigan et al. 2017; Pearce et al. 2017; T.K. Walsh et al. 2018). Importantly, the resistance toward pyrethroids in *H. armigera* is due to a unique P450 enzyme, *CYP337B3*, that is capable of metabolizing these pesticides into non-toxic hydro-pyrethroids (Joussen et al. 2012). *H. zea*, in contrast, remains susceptible to many major insecticides and tends to cause less damage than its Old World relative, although there are recent reports of resistance to *Bt*-toxins (Moar et al. 2010; Brévault et al. 2015; Reisig and Kurtz 2018). Therefore, one of the major concerns following the invasion of *H. armigera* is the formation of novel hybrid ecotypes or the introduction of genes associated with insecticide resistance and host-range expansion.

Anderson *et al.* (Anderson et al. 2018) unambiguously identified nine genetically intermediate individuals using whole-genome sequences between the two species from 2013, at least six to eight years after its possible initial introduction into Brazil, when significant crop damages were reported (Tay et al. 2013; Sosa-Gómez et al. 2016; Tay et al. 2017). Of the limited individuals analysed, eight were genetically closer to *H. armigera* with some introgressed haplotypes from *H. zea* scattered throughout the genome. The ninth individual resembled a F1 hybrid with stretches of homozygosity for each parental species, and all *H. armigera* surveyed individuals contained the *CYP337B3* gene involved in pesticide resistance. The apparent rarity of backcrossing to *H. zea* in these individuals (Anderson et al. 2018) initially suggested that hybridization would not introduce novel insecticide resistance to *H. zea* as had been feared (Mallet 2018). However, intensive pesticide-based control programs in Brazil aimed at controlling the invasive *H. armigera* provide a scenario in which there is strong selection for adaptive introgression of insecticide resistance genes into *H. zea* (Leite et al. 2014; Durigan et al. 2017; Fábio Pinto et al. 2017). These efforts to control *H. armigera* may also have had the converse effect of facilitating the establishment of *H. armigera* due to differential survival enabled by its resistance alleles (Norris et al. 2015; Dermauw et al. 2018).

Here, we use whole-genome resequencing of individuals in two temporal samples, soon after the outbreak of *H. armigera* in Brazil (2012-2013) and more recent populations (2016-2017) from a region with large areas cultivating cotton, soybean, and corn to examine the evolutionary outcomes following secondary contact between the *Helicoverpa* sibling species. We first identify genome-wide patterns of species distinctiveness and then explore signatures of introgression. We find that the species boundaries are strongly maintained in recent Brazilian populations. Then, we show that introgression is heterogeneous throughout the genome and that admixture has decreased over time in the Brazilian *H. armigera* population. Conversely, for *H. zea* there is a single strong signature of introgression around a region containing a novel, chimeric gene implicated in insecticide resistance (*CYP337B3* gene) which was previously only documented in *H. armigera* and identified in the invasive population in Brazil (Anderson et al. 2018; T.K. Walsh et al. 2018).

## Results

### Sympatric *H. zea* and *H. armigera* post-incursion persist as two well-distinct species

We analysed whole-genome sequence data from 137 moths from across the world: 90 *H. armigera armigera* (hereafter *H. armigera*), 39 *H. zea*, two hybrid individuals (hybrids considered as F1-F3), and six *H. punctigera*. The Brazilian individuals were collected in two temporal samples: soon after the *H. armigera* expansion in 2012-2013 (Sosa-Gómez et al. 2016) (59 individuals hereafter 2013) and more recent populations from 2016-2017 (37 individuals hereafter 2017). The Brazilian sampling included three regions of sympatry in Brazil that resulted from the invasion of *H. armigera* into local *H. zea* territory. We also included, as baseline comparison, four pre-invasion Brazilian *H. zea* individuals sampled in 2006 (Behere et al. 2007), as well as eight individuals of *H. zea* from allopatric (USA) populations. Additionally, nine individuals of *H. armigera* from the Old World which are all considered to be geographically or temporally isolated from the Brazilian invasion (Fig. 1A, Table S1-3), as well as 14 *H. a. conferta* individuals from Australia, and six individuals from an outgroup Australian endemic species *H. punctigera* (Fig. 1A).

**Fig. 1.**
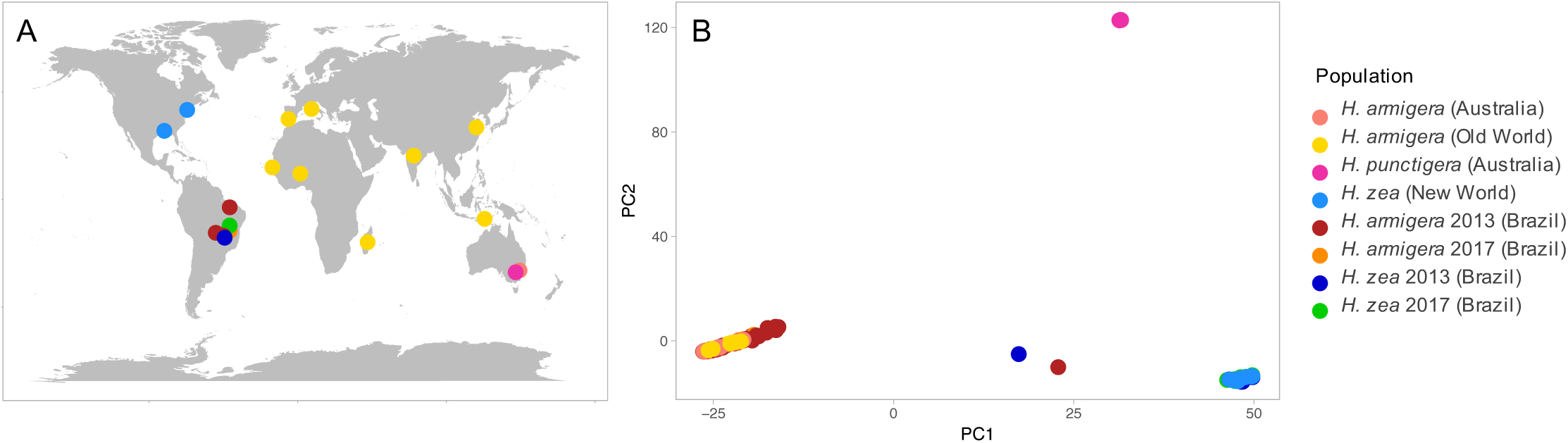
**Sampling locations and Principal Component Analysis. A**. Sampling locations of *Helicoverpa* individuals. **B.** Genetic PC1 and PC2 of 50,000 genome-wide SNPs. Colour codes indicate different samples and species based on mitochondrial sequences, and the groups used for the analyses. *H. armigera* (Australia) refers to the subspecies *H. armigera conferta* and *H. armigera* (Old World) to the subspecies *H. armigera armigera* in the rest of the Old World. *H. zea* (New World) comprises Brazilian pre-invasion and USA, and finally *H. armigera* 2013/2017 and *H. zea* 2013/2017 refer to the post-invasion sympatric populations in Brazil.

Principal component analysis (PCA) based on 50,000 whole-genome single-nucleotide polymorphism data separated *H. zea* from *H. armigera* on principal component 1 which explained 17.2% of the variation in the data (Fig. 1B). Across the wide geographic range covered by our sampling there was no evidence of population substructure. The reported separation between *H. armigera conferta* from Australia and *H. armigera armigera* from the rest of the Old World (Anderson et al. 2016; Anderson et al. 2018) was evident when resuming at least the first 25 principal component axes (Fig. S1). Two individuals fell between *H. armigera* and *H. zea,* likely representing early generation hybrids. The remaining 97 individuals from Brazil clearly have a closer genetic affinity to one or other of the parental species. Consistent with PC1, ADMIXTURE (Anon) analyses showed strong differentiation between the two species, which is supported by the lowest cross-validation error (CV) of any assessed when the number of populations is K=2 (Fig. S2).

To classify parental and hybrid genotypes, we calculated the proportion of ancestry and heterozygosity based on SNPs that strongly segregated between allopatric populations of the species. In accordance with the PCA and ADMIXTURE results, the classification of Brazilian individuals identified only two early generation hybrids (Fig. 2). One individual resembled an F2 hybrid from 2013, while the other individual resembling F3 hybrids from 2013 and 2017, respectively. All remaining individuals grouped with one of the parental species, although with various fractions of their genome coming from the other species (Proportions of ancestry listed in Table S4). A total of 66 individuals were closer to *H. armigera* with ∼ 0 - 9.2% of alleles derived from *H. zea*. Meanwhile, the 27 individuals that grouped with *H. zea* samples had 0 - 4.1% of *H. armigera* ancestry (Table S4). The tight within-species clustering and low prevalence of intermediate individuals indicated that almost none of the recently sampled individuals resulted from contemporary hybridisation, suggesting that hybridisation is rare on a per-individual basis in recent populations.

**Fig. 2.**
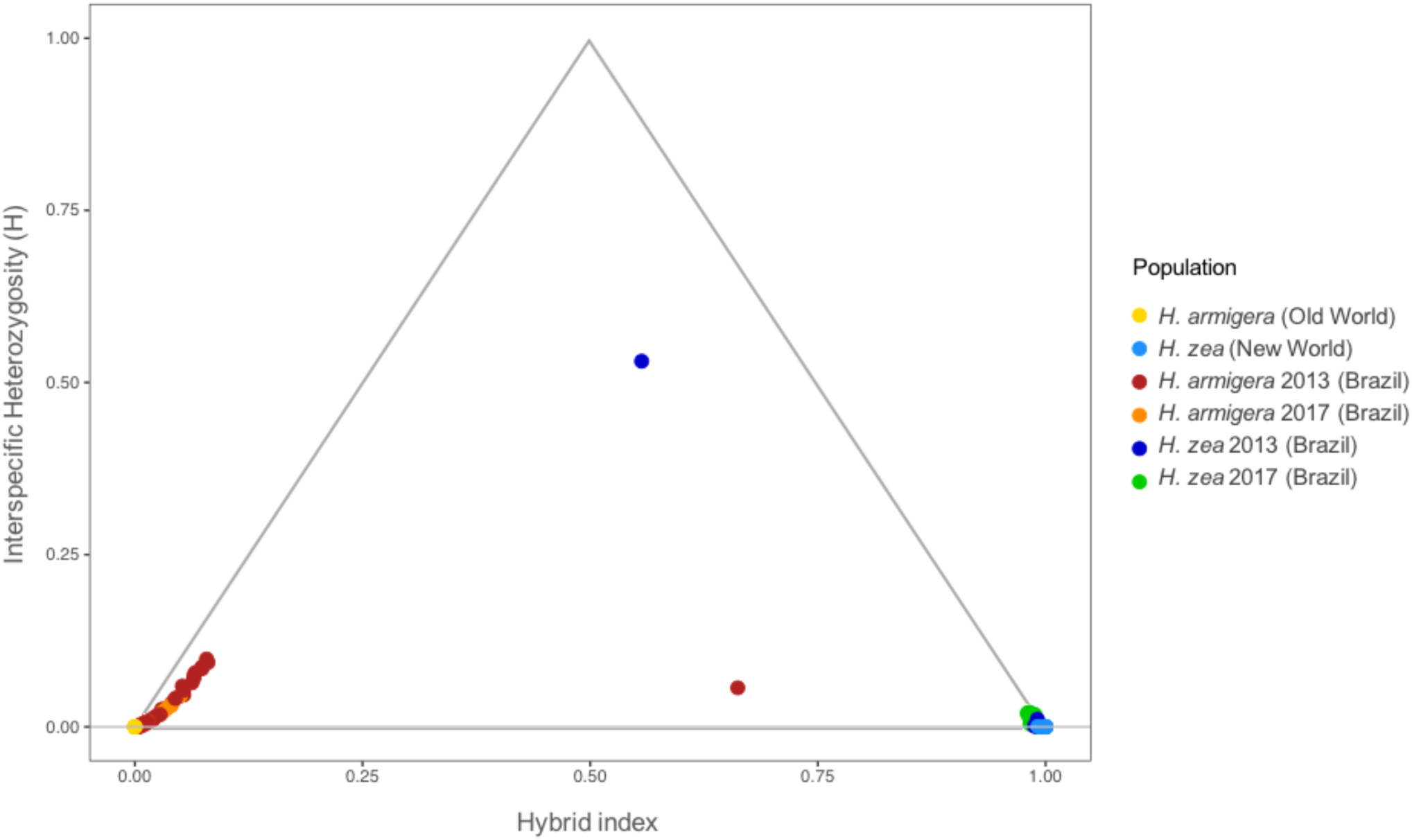
**Classification of hybrid and parental genotypes**. Classification is based on the hybrid index and interspecific heterozygosity for *H. zea* and *H. armigera* individuals. The hybrid index and the interspecific heterozygosity were calculated based on ∼1,000,000 SNPs that are fixed differences between allopatric populations of the parental species.

### Reduced interspecific divergence in sympatry around a single genomic region

We characterized patterns of divergence between sympatric and allopatric pairs of *H. zea* and *H. armigera* populations using the fixation index *F_ST_*. We found similar high genome-wide differentiation between *H. zea* and *H. armigera* for allopatric and recent sympatric populations with a mean autosome-wide *F_ST_* of 0.474±0.001 and 0.496±0.001, respectively (Fig. S3, Table S1). By contrast, *F_ST_* values for intraspecific comparisons were low, consistent with a clear genetic identity within species (Fig. S3).

To compare differentiation across the genome, we calculated *F_ST_*-values in 100-kb non-overlapping windows. Interspecific *F_ST_*-values for allopatric and sympatric populations were consistently high throughout the genome except for a single region on chromosome 15 (Fig. 3C-F). This region included the insecticide-resistance gene *CYP337B3*, whereby *F_ST_* dropped to 0.14 for the sympatric Brazilian populations of *H. armigera* and *H. zea* (Fig. 3C, E). Intraspecific *F_ST_*-values across the genome between Brazilian and Old World *H. armigera* populations were also consistently low, although there was a slight peak at the *CYP337B3* locus (Fig. S3B). Between *H. zea* from USA and Brazil, there was also only one peak of differentiation, corresponding to the same region on chromosome 15 which contains the insecticide-resistant locus and that showed low differentiation for the interspecific comparison between Brazilian *H. zea* and *H. armigera* (Fig. 3D). Although both *F_ST_* estimates at a genome-wide scale and across windows show that differentiation between species is virtually the same for sympatric and allopatric populations, these patterns of reduced between-species and elevated within-species differentiation around the same region, hint at recent localized introgression.

**Fig. 3.**
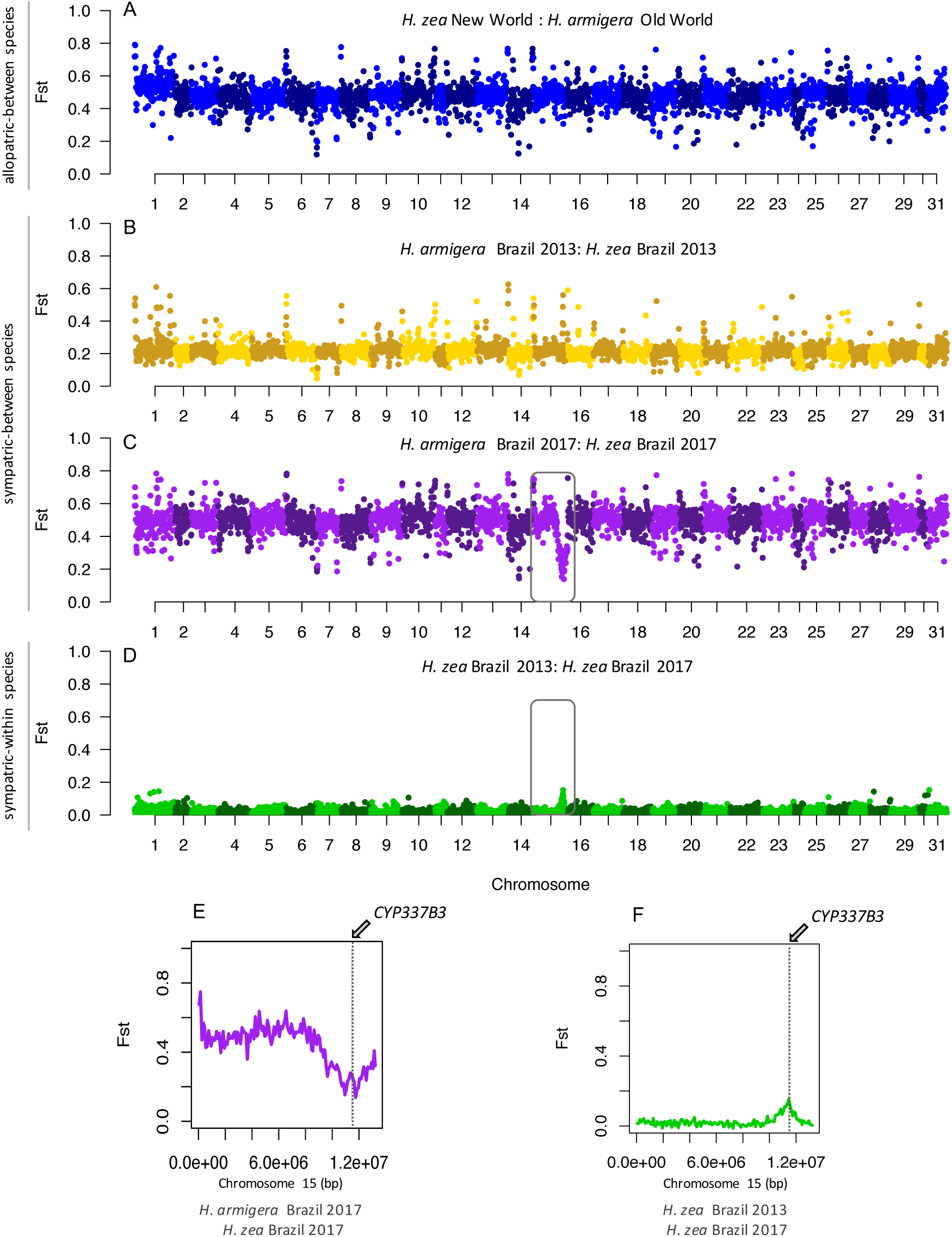
***F_ST_* scans across the genome.** *F_ST_* scans across 100kb genomic windows between sympatric and allopatric populations of *H. armigera* and *H. zea.* Squares on panels **C** and **D** highlight chromosome 15 and panels **E** and **F** zoom in on *F_ST_* values on chromosome 15 showing the location of the *CYP337B3* gene. Indeed, the *F_ST_* between the invasive Brazilian and Old World *H. armigera* was 0.0062±0.0001 (Fig. S3A), and the pairwise *F_ST_* between Brazilian *H. zea* and North American *H. zea* averaged 0.0118±0.0002 (Fig. S3A).

### Widespread phylogenetic discordance is consistent with recent gene flow

We explored species phylogenies across the genome using Twisst (Martin and Van Belleghem 2017), which quantifies the frequency of alternative topological relationships among all individuals in sliding windows of 50 SNPs. These phylogenetic comparisons across the genome indicated large-scale introgression among recent samples of Brazilian *H. armigera* and *H. zea* (Fig. 4). The three possible unrooted topologies that describe the relationships between allopatric and sympatric populations were represented across the genome (Fig. S4). Around 60% of the windows had completely sorted genealogies (i.e. weighting of 1), most of which matched the expected species branching order (Blue in Fig. 4, Fig. S4). Around 90% of the topologies are concordant with the species tree (Fig. S4). The two discordant topologies are likely explained by incomplete lineage sorting of the non-structured populations of *H. zea* and *H. armigera* (Green and orange in Fig. S4). However, the topology that groups recent *H. zea* and *H. armigera* together from Brazil is more common and is consistent with a recent burst of gene-flow between sympatric species and introgression (Green in Fig. 4, Fig. S4). Notably, from these regions of phylogenetic incongruence, the longest continuous interval with a discordant topology was on chromosome 15, around the *CYP337B3* locus (Fig 4). Therefore, given that introgression seems to be restricted to a single region, the similar frequencies of the non-species phylogenies suggest that introgression in recent populations of *H. zea* is not widespread.

**Fig. 4.**
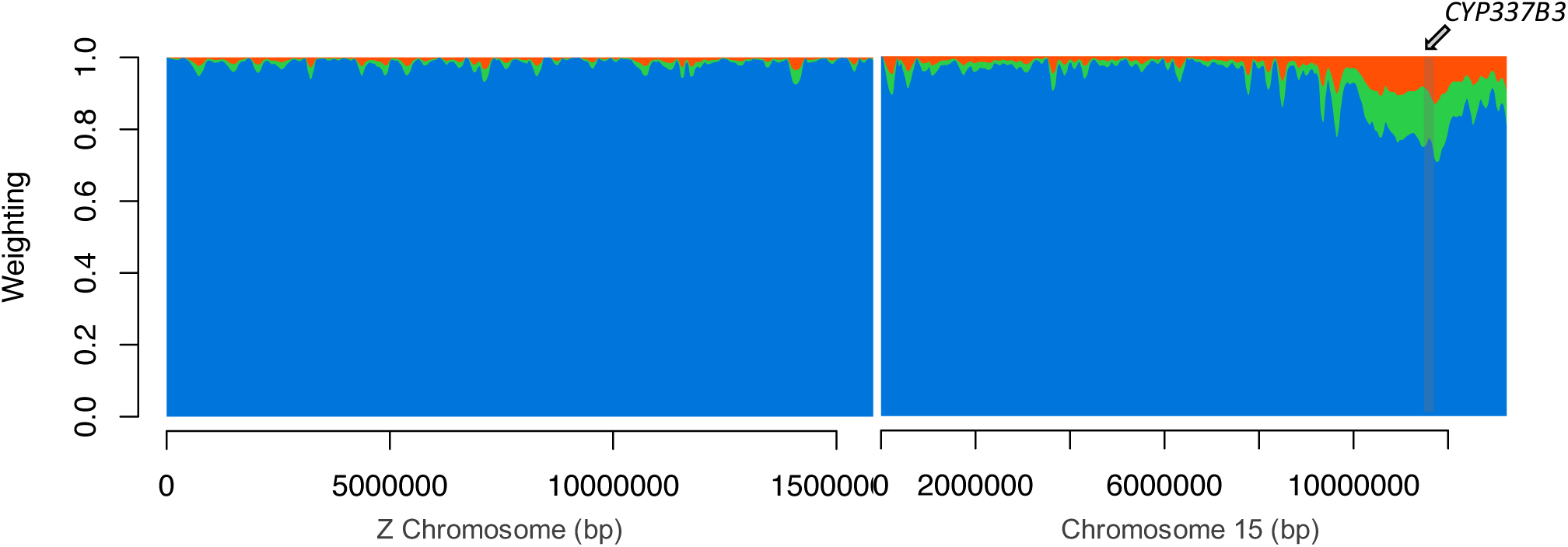
**Topology weightings across the Z-chromosome and chromosome 15.** Twisst analysis on 50 SNPs windows for a species topology with the relations: ((Old World *H. armigera*, Brazilian *H. armigera* 2017) (New World *H. zea*, Brazilian *H. zea* 2017)). Blue represents the weight of the species topology ((Old World *H. armigera*, Brazilian *H. armigera* 2017) (New World *H. zea*, Brazilian *H. zea* 2017)). Green ((Brazilian *H. zea* 2017, Brazilian *H. armigera* 2017) (New World *H. zea*, Old World *H. armigera*)) and orange ((Brazilian *H. zea* 2017, Old World *H. armigera*) (New World *H. zea*, Brazilian *H. armigera* 2017)), are alternative topologies to the species tree. See Fig. S4 for explanations of the topologies.

### Introgression is biased to specific genomic regions and has decreased over time

To test for genetic admixture, we calculated Patterson’s D statistics which test for an imbalance of discordant SNPs of derived alleles which is indicative of introgression (Green et al. 2010; Patterson et al. 2012). We found strong signals of gene-flow between Brazilian *H. zea* and *H. armigera* for 2013 and 2017 samples (Table 1). We then examined *f*, the proportion of the genome that has been shared between species (Green et al. 2010), and observed a strong trend to a decreased *f* over time in *H. armigera* (Fig. 5, Table S2). The fraction of ancestry derived from their sibling species was higher in *H. armigera* populations than for *H. zea* in 2013, suggesting asymmetric introgression early after the invasion (Fig. 5).

**Fig. 5.**
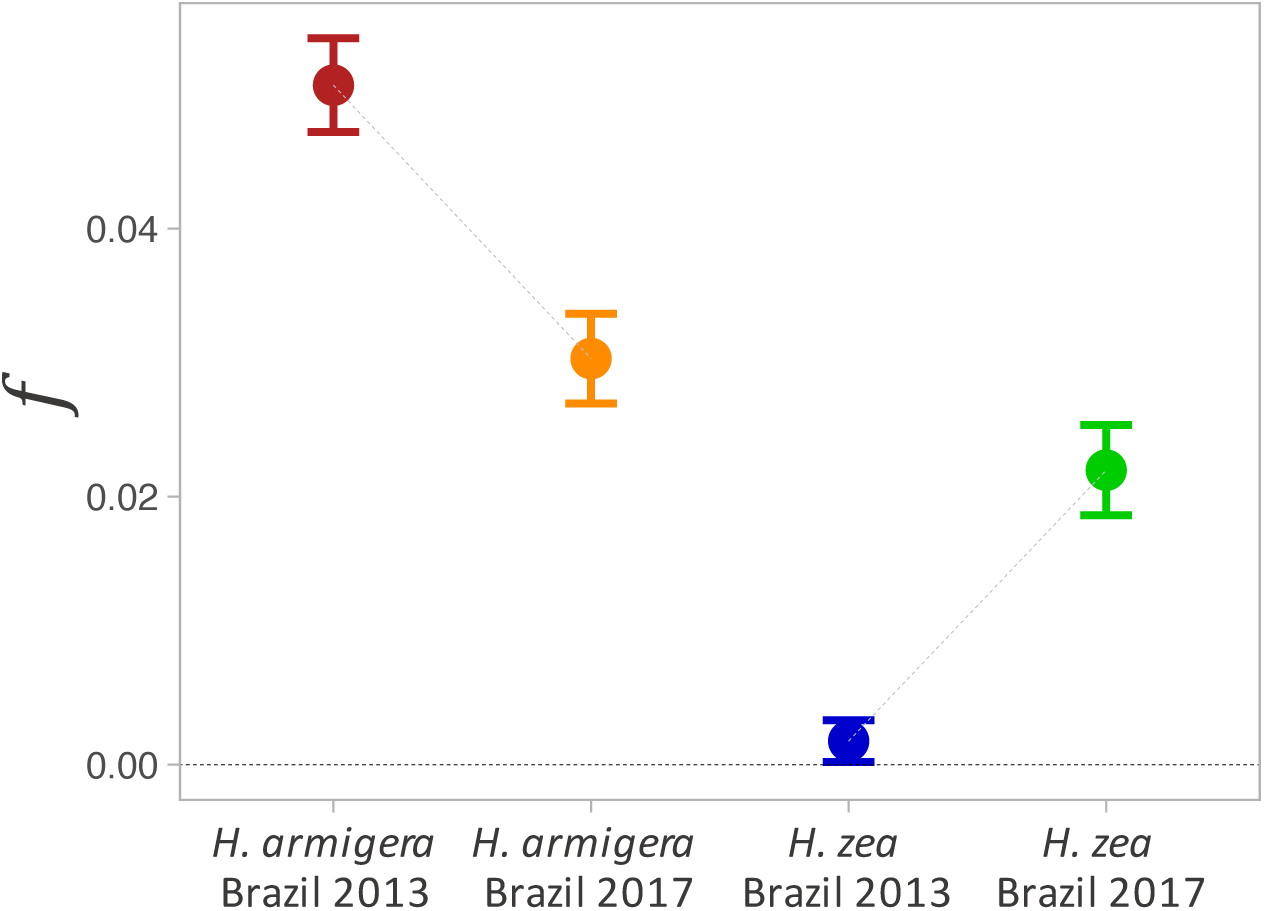
**Genome-wide proportion of admixture.** Proportion of admixture (*f*) estimated over time for *H. armigera* and *H. zea* in Brazil estimated as an F4 ratio for the genealogy ((Australian *H. armigera conferta,* Old World *H. armigera*; X; New World *H. zea*).

**Table 1.** **Results of D-statistic.** Admixture test for Brazilian populations of *Helicoverpa* in 2013 and 2017.

| Target population | D-statistic | Z-score | P-value |
| --- | --- | --- | --- |
| *H. armigera* 2013<sup>a</sup> | 0.1203 ± 0.005015 | 23.986 | < 0.0001 |
| *H. armigera* 2017<sup>a</sup> | 0.0259 ± 0.004774 | 5.427 | < 0.0001 |
| *H. zea* 2013<sup>b</sup> | 0.0068 ± 0.004373 | 1.551 | 0.1209 |
| *H. zea* 2017<sup>b</sup> | 0.0554 ± 0.010163 | 5.455 | < 0.0001 |

Model used for *H. armigera* populations ((A,B)C)D): (((x, *H. armigera* Old World) *H. zea* New World) *H. punctigera*).
Model used for *H. zea* populations ((A,B)C)D): (((x, *H. zea* New World) *H. armigera* Old World) *H. punctigera*). A significantly positive Z-score implies gene flow between A and C. A significantly negative Z-score implies gene-flow between B and C. P-values <0.05 indicates D-statistic significantly different from 0.

To locate introgressed loci, we used the summary statistic *f_d_*, since it is better suited than *f* to quantify admixture in small genomic regions and which is roughly proportional to the effective migration rate (Martin et al. 2015). As expected from the genome-wide *f*, *f_d_*-values also decreased over time in populations of sympatric *H. armigera* and *H. zea* (Fig. 6). However, the proportion of admixture estimated in sliding windows showed considerable heterogeneity in the extent of introgression across the genome (Fig. 6). In both temporal samples and species, admixture was minimal on the Z-chromosome, particularly in *H. armigera,* indicating a strong barrier to introgression (Fig. 6B, D). For Brazilian *H. armigera* we found high heterogeneity in admixture proportion in the autosomes, with peaks scattered throughout different chromosomes (Fig. 6D). Some regions, including the entire sequence of chromosome 15, exhibited *f_d_-*values below the genome-wide average, implying strong localised barriers to gene-flow. Conversely, in local *H. zea* populations, *f_d_* values were lower throughout the genome compared to the invasive *H. armigera* in 2013. Yet, for recent *H. zea* populations, we found a peak showing a higher proportion of admixture (Fig. 6B), than any peak in *H. armigera* and reflecting almost complete replacement. This peak was located across the region on chromosome 15 containing the *CYP337B3* gene responsible for pyrethroid resistance and that initially was only present in the invasive species (Fig. 6). Additionally, gene scans of *H. zea* revealed that *CYP337B3* was present in 14 out of 18 *H. zea* individuals from 2017 (frequency ∼0.70) (Table S4) and that the sequence was identical to that found in the invasive *H. armigera* population.

**Fig. 6.**
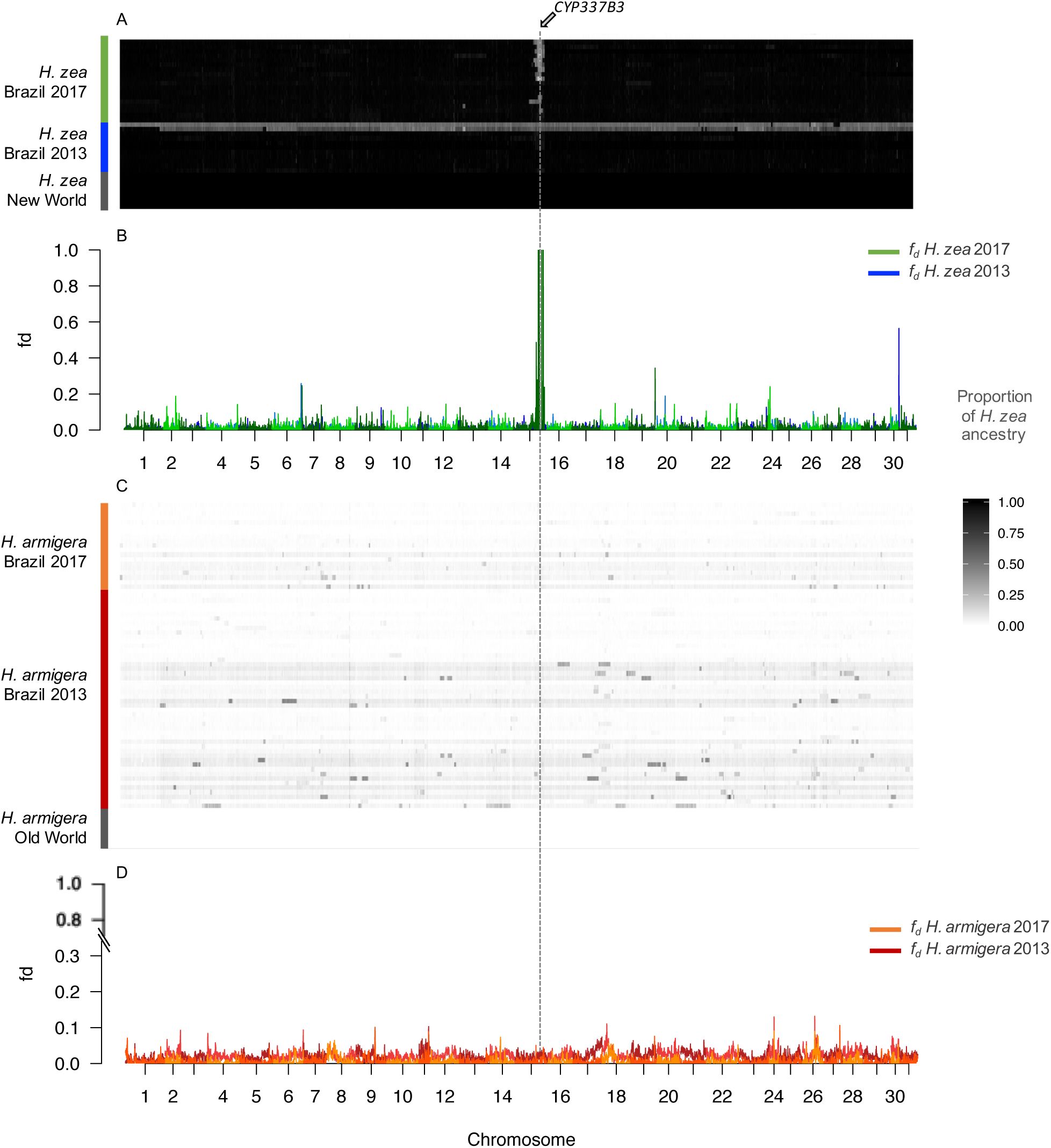
**Proportion of admixture across the genome.** (*f_d_*) and introgressed tracks per individual. **A.** Admixture tracks displaying *H. zea* ancestry in 100kb windows for individuals of *H. zea* 2013, *H. zea* 2017, and *H. zea* (New World). **B.** Proportion of admixture *f_d_* across 100kb windows, for populations of *H. zea* 2013 (blue) and *H. zea* 2017 (green). **C.** Admixture tracks displaying *H. zea* ancestry in 100kb windows for individuals of *H. armigera* 2013, *H. armigera* 2017, and *H. armigera* (Old World). **B.** Proportion of admixture *f_d_* across 100kb windows, for populations of *H. armigera* 2013 (dark red) and *H. armigera* 2017 (orange).

Increased *F_ST_* values in both within-species comparisons around the *CYP337B3* locus might also be a result of within species population dynamics. However, the joint estimation of *F_ST_* and *f_d_* allowed us to disentangle linked selection in *H. armigera* from differentiation due to admixture in *H. zea. F_ST_* is expected to be highly heterogeneous even in the absence of gene flow, with elevated values in regions of the genome that experience strong linked selection (Jakobsson et al. 2013; Martin et al. 2015). Therefore, the weak *F_ST_* peak within *H. armigera* is likely an artefact of the recent selective sweep in this region in chromosome 15 (T.K. Walsh et al. 2018) (Fig. S3). In contrast, the *F_ST_* peak on chromosome 15 in the within-species comparison of *H. zea* populations pre- and post-*H. armigera* invasion coincided with the *f_d_* peak in recent *H. zea* populations, implying that differentiation was caused by introgression.

We then calculated the proportion of *H. zea* ancestry across the genome of all individuals using a data set of diagnostic SNPs that differentiate the two species and estimated a hybrid index in 100-kb sliding windows (Fig. 6A). Across individuals, there was evidence for long regions acquired through introgression (Fig. 6A). Consistent with the *f_d_* analyses, recent individuals of Brazilian *H. zea* exhibited a single clearly introgressed block on chromosome 15, around the *CYP337B3* locus.

By contrast, in early *H. armigera*, long introgressed regions were pervasive, indicating introgression of almost whole chromosomes. Particularly long admixture tracks derived from *H. zea* on chromosome 18 seemed to be persistent across most *H. armigera* individuals in 2013 and 2017. Nevertheless, we observed a general pattern of shorter introgressed blocks and a lower effective migration rate in recent populations of *H. armigera*, implying that admixed regions have been substantially removed and broken up by recombination over time.

To characterize recent selection around the *CYP337B3* locus we calculated Tajima’s D on 10-kb windows across the region on chromosome 15 containing this gene. There was a trough in Tajima’s D consistent with a selective sweep at the location of *CYP337B3* from invasive *H. armigera* in 2013 (Fig. 7A, Tab S3) that is absent in the non-introgressed *H. zea* from the same period (Fig. 7B, S5), thus suggesting strong selection on this locus. In recent populations of Brazilian *H. armigera*, Tajima’s D is negative in the vicinity of the insecticide resistance allele (Tajima’s D = −0.60) although not lower compared to the genome-wide average (Tajima’s D ± sd: −0.98±0.36) (Fig. 7A). In contrast, recent populations of *H. zea* harbouring the *CYP337B3* haplotype exhibited significantly lower Tajima’s D along this region (Tajima’s D = −0.72; genome-wide Tajima’s D ± sd: 0.57±0.59) suggesting recent strong selection on this introgressed locus (Fig. 7B, Tab S3).

**Figure 7.**
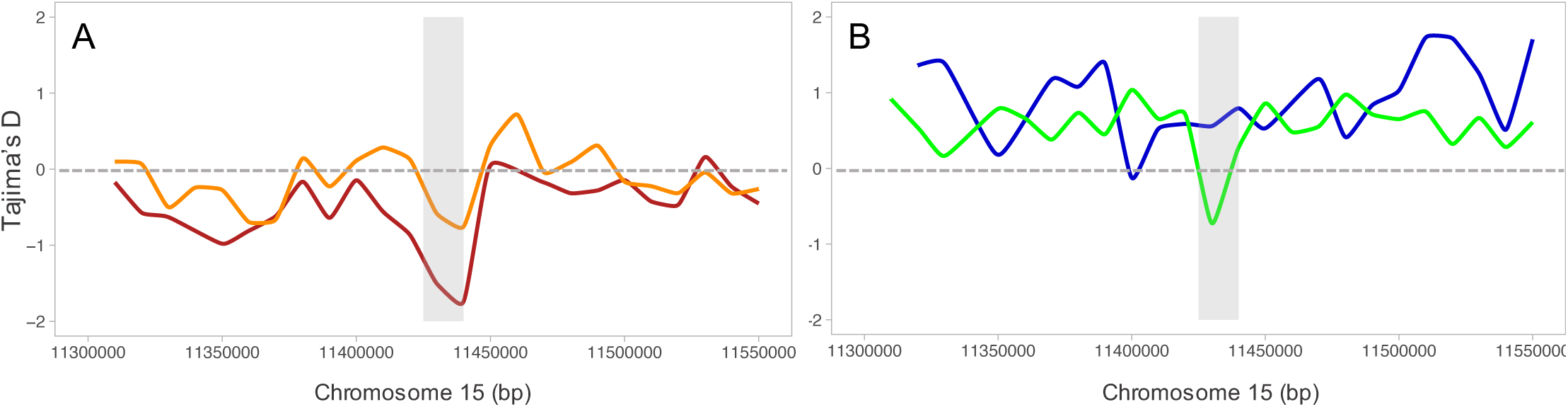
Tajima’s D in 10kb windows along the region in chromosome 15 containing *CYP337B* gene for populations of *H. armigera* and *H. zea* in Brazil. **A.** Tajima’s D for *H. armigera* 2013 (dark red) and for *H. armigera* 2017 (orange). **B.** Tajima’s D for *H. zea* 2013 (blue) and for *H. zea* 2017 (green) containing *CYP337B3* gene. Grey shading indicates the location of *CYP337B3*.

## Discussion

We have used whole genome sequencing of 137 individuals to document the interaction between native and invasive crop pest species. This has demonstrated that these species are maintaining their differentiation in the face of hybridisation, showing evidence for a reduction in admixture over time. However, a single locus associated with insecticide resistance shows dramatic evidence for an introgressed selective sweep from *H. armigera* into *H. zea*. Whole genome sequencing therefore offers an opportunity to track the fate of invasive species in the face of hybridisation. In this particular case, we are witnessing the emergence of a new paradigm where insecticide resistance or other adaptive traits can be passed rapidly between species in the native range and might eventually lead to the creation of a megapest species. This has serious implications for pest control and agricultural crop production in South America and more broadly around the world as movement of pests is not unidirectional.

### Sympatric *H. zea* and *H. armigera* persist as two well-differentiated species

In secondary contact after strict allopatric speciation, there are a number of possible outcomes, with barriers to gene flow potentially being either strengthened or broken down and resulting in the fusion of the divergent species. Despite pervasive introgression following invasion and secondary contact between *H. armigera* and *H. zea*, we found that the species boundaries are strongly maintained. Most individuals showed some proportion of introgression, but early hybrids and backcrosses were scarce, suggesting low levels of hybridisation. Differentiation between species is roughly the same between allopatric and sympatric populations except for a single “pocket” of introgression driven by strong selection. Therefore, from the myriad of possible outcomes after secondary contact, our results do not lend support to *H. armigera* and *H. zea* fusing back into one species, nor formation of a new hybrid species, at least not in the time frame examined here. On the contrary, the predominance of parental forms and the rarity of hybrids particularly in the recent samples suggest strong selection against introgression.

The *Helicoverpa* sibling species exhibit both prezygotic and postzygotic barriers that can account for the low rates of natural hybridisation (Hardwick 1965; Laster and Sheng 1995; Berg et al. 2014; Mallet 2018). Despite experiments having demonstrated no instances of sterility in *H. armigera* x *H. zea* hybrids in reciprocal crosses inbred for two generations or in lines backcrossed for four, they exhibit strong assortative mating (Hardwick 1965; Laster and Sheng 1995). Mating is particularly difficult to obtain between F1 hybrid and *H. zea* (AZ x Z), which can account for the lower admixture proportion we observe in *H. zea* populations in 2013. Traits linked to prezygotic isolation include morphological differences in the male and female genitalia as well as divergent pheromonal bouquets (Hardwick 1965; Laster and Sheng 1995; Berg et al. 2014). A possible outcome of mating between *H. zea* and *H. armigera* is to ‘lock up’, leading to death without reproduction of the individuals involved (Hardwick 1965). This is apparently due to differences in the length of an elaborate inflating appendix in the genitalia, which makes the male unable to successfully withdraw his aedeagus extension from the female (Mallet 2018). Although this can also occur in intraspecific mates, this clogging is more common in interspecific matings (Hardwick 1965; Mallet 2018). Furthermore, behavioural tests in *Helicoverpa* moths showed that female pheromones and their ratios are used as critical signals to avoid interspecific matings (Fadamiro and Baker 1997; Quero et al. 2001; Berg et al. 2014). Even though *H. zea* and *H. armigera* both use Z-11-hexadecenal and Z-9-hexadecenal as the primary sex pheromones emitted by females and detected by males, the ratio of the latter to the former is about 3-fold higher in *H. armigera* (Fadamiro and Baker 1997; Quero and Baker 1999; Quero et al. 2001; Berg et al. 2014). Therefore, both mechanical and chemical reproductive isolation likely play a role in assortative mating and the reduction of successful interspecific matings.

In addition to pre-zygotic isolation, selection against hybridization could be the result of intrinsic postzygotic barriers including Bateson-Dobzhansky-Müller incompatibilities (BDMIs) or alternatively “pathway” incompatibilities. BDMIs cause reduced fitness in hybrids but can be broken down by recombination in backcross progeny (Bomblies et al. 2007; Lee et al. 2008; Schumer et al. 2018). Conversely, “pathway” incompatibilities reduce fitness in recombinant hybrids in which co-adapted alleles become separated (Lindtke and Buerkle 2015). Simulations and empirical data show that BDMIs usually have an even, genome-wide effect whereas “pathway” incompatibilities produce distinct localised barriers to introgression (Lindtke and Buerkle 2015; Martin et al. 2019). Our results showed a genome-wide reduction of introgression over time, thus suggesting pervasive selection against hybridization likely as a result of widespread BDMIs throughout the genome. Moreover, patterns of differentiation captured by *F_ST_* also suggested highly polygenic barriers that maintain *H. armigera* and *H. zea* as distinctive species despite the recent burst in gene flow. BDMIs are more likely to act as polygenic barriers and are a central mechanism underlying reproductive isolation of species formed through allopatric divergence as in the case of *H. armigera* and *H. zea* (Bomblies et al. 2007; Lee et al. 2008). Therefore, we hypothesize that barrier loci like BDMIs accumulated rapidly in the 1.5 My of allopatric divergence and once in secondary contact these incompatibilities contribute to selection against admixture across the genome. In summary, our findings highlight that species can be unambiguously different but coupled by the interchange of adaptive genes.

### Rapid adaptive introgression of an insecticide resistance gene

Several lines of evidence suggested rapid adaptive introgression between the *Helicoverpa* sibling species after the *H. armigera* invasion in South America. In order to provide evidence for adaptive introgression, the introgressed haplotype needs to be shown to be responsible for a phenotype with a beneficial impact on fitness, and to have increased in frequency under the action of selection. The candidate region containing *CYP337B3* appears to satisfy both requirements. First, pesticide resistance toward pyrethroids is due to the unique P450 enzyme encoded by this gene (Joussen et al. 2012; T.K. Walsh et al. 2018). This chimeric enzyme arose independently at least eight times from unequal crossover events between two parental P450 genes (T.K. Walsh et al. 2018), and is capable of metabolizing pyrethroids to non-toxic compounds in larvae and adults (Joussen et al. 2012). Indeed, the exclusive presence of *CYP337B3* has been shown to confer a 42-fold resistance to fenvalerate in *H. armigera* (Joussen et al. 2012). Furthermore, Durigan *et al.* (Durigan et al. 2017) showed that *CYP337B3* is associated with the very low mortality of larvae when sprayed with the pyrethroids deltamethrin and fenvalerate in invasive *H. armigera armigera* in Brazil.

Second, the *CYP337B3* introgression into *H. zea* and subsequent increase in frequency coincided with an intensification in pesticide-use in Brazil directed to control outbreaks generated by the invasive *H. armigera* (Leite et al. 2014; Durigan et al. 2017; Fábio Pinto et al. 2017). In the 2012/2013 growing season, cotton, soybean, and maize crops were impacted by *Helicoverpa* species at unusually high infestations resulting in billion dollar losses to Brazilian agriculture (Durigan et al. 2017). Initially, the species was presumed to be *H. zea,* but during this season, growers reported a reduced efficacy of insecticides despite the increase in sprays and the known susceptibility of *H. zea* to pesticides (Durigan et al. 2017). The resistant individuals were subsequently confirmed as *H. armigera*, showing it to have successfully spread and become established after being introduced into Brazil from approximately 2007 onwards (Tay et al. 2013; Lopes-da-Silva et al. 2014; Tay et al. 2015; Sosa-Gómez et al. 2016). Furthermore, the survival and establishment of *H. armigera* in the studied region during the 2012/2013 season may have been facilitated by a decline in the use of Bt cotton cultivars during this season due to problems with the quality of fibres produced. Since *H. armigera* was detected in 2013, the frequency and dosages of insecticide sprayings including pyrethroids dramatically increased (Durigan et al. 2017). Thus, the survival of early hybrids and backcrosses enabling introgression is likely related to the rapid scale-up of insecticide-based control measures in Brazil after the outbreak generated by the expansion of *H. armigera*. The observation of early generation hybrids mostly during the outbreak in 2013 and not in recent samples, as well as the selective sweep in early *H. armigera* around *CYP337B3* further supports this hypothesis. The hypothesis of recent selection on this locus is supported by our population genomic analysis demonstrating a signature of recent selection at the introgressed allele in *H. zea* that is not present in the background of the ancestral allele. Meanwhile, *H. zea* populations which were susceptible to pyrethroids, seem to have responded to insecticide selection pressure by co-opting resistance from *H. armigera* through adaptive introgression.

The introgression of a gene involved in insecticide resistance from *H. armigera* into *H. zea* joins a series of animal examples of rapid adaptive introgression triggered by human-mediated alteration of fitness landscapes. These examples include the European house mouse, *Simulium* black flies, and *Anopheles* mosquitos whereby the increase in pesticide exposure acted as a selective force sufficient to drive introgression across species boundaries (Djogbénou et al. 2008; Adler et al. 2010; Song et al. 2011; Clarkson et al. 2014; Norris et al. 2015). Remarkably, in all these cases adaptive introgression occurred from the species with a higher effective population size into the species with lower *Ne* and lower genetic diversity. This pattern has also been found in natural populations whereby the selective agent is not human-mediated. For instance, adaptive introgression of colour pattern genes from *Heliconius melpomene* into *H. timareta* because of natural selection for mimicry (Pardo-Diaz et al. 2012; Edelman et al. 2019), tidal marsh adaptations from *Ammospiza caudacauta* into *A. nelson* sparrow (J. Walsh et al. 2018), and female-competitive traits from *Jacana spinose* into *J. jacana* (Lipshutz et al. 2019). This pattern is expected given that species with higher *Ne* harbour more standing variation which is the raw material for adaptation and introgression is thus less likely to add beneficial variants (Barton 2010; Jensen and Bachtrog 2011). Moreover, small populations have a higher mutational load because they are more prone to accumulate deleterious mutations whereas large populations are less subjected to drift (Kimura and Ohta 1969; Barton 2010). However, the most famous example of adaptive introgression from Neanderthals and Denisovans into out-of-Africa human populations seems to be in the reverse direction, undermining the generality of this pattern and highlighting the bidirectional nature of introgression (Green et al. 2010; Patterson et al. 2012; Gazave et al. 2014; Huerta-Sánchez et al. 2014; Mafessoni and Prüfer 2017). Of note, Neanderthals were established in Eurasia when humans were expanding into Europe, and although less is known about introgression from humans to Neanderthals and Denisovans, there is evidence of gene flow in this direction (Green et al. 2010; Patterson et al. 2012; Huerta-Sánchez et al. 2014; Prüfer et al. 2014).

The most comprehensive genomic study of rapid adaptive introgression resulting from pesticide pressure is between *Anopheles gambiae* and *A. coluzzi,* the major malaria vectors in Africa (Clarkson et al. 2014; Fontaine et al. 2015; Norris et al. 2015). Several studies have shown introgression of mutations conferring resistance to pyrethroids from *A. gambiae* into *A. coluzzi* (Djogbénou et al. 2008; Clarkson et al. 2014; Norris et al. 2015). The *CYP337B3* trickle from *H. armigera* into *H. zea* coincided with increased in pesticide spraying, similar to introgression of insecticide resistance alleles in *Anopheles* overlapping with the start of a significant insecticide-treated bed net distribution campaign in Mali (Norris et al. 2015). After introgression of the insecticide-resistant genes, these alleles almost reached fixation in four years in *A. coluzzi* populations (Norris et al. 2015) which is a temporal scale that strikingly resembles introgression and near fixation of the *CYP337B3* loci in *H. zea* in Brazil.

Despite the similitudes between *Anopheles* and *Helicoverpa* examples of rapid adaptive introgression, a critical difference relies on the speciation mechanism underlying the divergence between the sibling species. *A. gambiae* and *A. coluzzi* diversified in sympatry only about 400,000 years ago (Clarkson et al. 2014; Fontaine et al. 2015) whereas *H. armigera* and *H. zea* differentiated in strict allopatry for 1.5 Mya and came into contact only in the recent 2013 invasion (Pearce et al. 2017; Anderson et al. 2018). Accordingly, the genomes of *A. gambiae* and *A. coluzzi* exhibit very low differentiation except for two large regions that show exceptional divergence known as “genomic islands” of diversification (Clarkson et al. 2014). The insecticide-resistant alleles are located within one of these major diverged regions, and its introgression resulted in the homogenization of the entire island, though no apparent impact on reproductive isolation (Clarkson et al. 2014). Conversely, in *H. armigera* and *H. zea* the differentiation is not restricted to “islands” but rather we observed consistently high *F_ST_* values throughout the genome. Furthermore, incompatibilities are more likely to accumulate in allopatry, and therefore are expected to be more pervasive in *Helicoverpa* than in *Anopheles* sibling species (Bank et al. 2012). This suggests that in the face of intense selective pressure, adaptive variation can rapidly cross species boundaries even when differentiation and incompatibilities are pervasive across the genome, and not only between slightly differentiated species as in *Anopheles* mosquitos.

### Asymmetric introgression at the front of an invasion

We have primarily focused on the adaptive introgression of *CYP337B3* from the invasive *H. armigera* into local *H. zea* even though we observed a higher effective migration of alleles from *H. zea* into *H. armigera.* This pattern is in agreement with theoretical models of secondary contact after invasions that account for demographic processes, which predict greater introgression of neutral genes from the local into the foreign species (Currat et al. 2008; Mesgaran et al. 2016; Quilodrán et al. 2019). This asymmetry is partly explained by a demographic imbalance between the two species at the wave front, where the invading species is at lower densities (Currat et al. 2008; Hall 2016; Mesgaran et al. 2016). Therefore, it is likely that early invading *H. armigera* individuals had few available conspecific mates, and thus, were involved in more heterospecific crosses.

Most of the admixed alleles in the invasive *H. armigera* are likely to be neutral (Currat et al. 2008). However, introgression can also provide variants that have already been tested by natural selection and in precisely the geographic area in which they are already adapted (Martin and Jiggins 2017). Both models and empirical data show that genes from the resident species associated with local adaptation are easily introgressed into the invading species (Currat et al. 2008; Saarman and Pogson 2015). Contrastingly, genes from the invading species introgress into the local species only if they are under very strong positive selection in the local environment (Currat et al. 2008), as appears to be the case for the *CYP337B3* gene. For genes with a large effect on fitness and a clear signal of admixture localized into a single region, it is reasonable to suggest that the increased frequency of the introgressed haplotypes was driven by strong positive selection. However, if introgression is pervasive, it is challenging to distinguish introgressed alleles that are beneficial from neutral genes that have also introgressed. While *H. zea* has developed resistance to two-toxin Bt maize and cotton (Cry1A and Cry2a) in North America (Reisig and Kurtz 2018), no single specific genes or mutations have yet been identified. Therefore, there are currently no plausible candidates for adaptive introgression from *H. zea* into *H. armigera* nor has any Bt resistance been observed in *H. zea* in South America. Nonetheless, studies on gene flow between *H. zea* populations across North, Central and South America would be valuable in understanding possible movement of resistance genes such as the *CYP337B3* gene introgressed from *H. armigera.* Further temporal monitoring of the invasive *H. armigera* populations in Brazil will permit the detection of fine-scale selection on any introgressed alleles that may be identified.

### Practical implications of rapid evolution triggered by invasive hybridization

The pervasive introgression between local *H. zea* and invasive *H. armigera* demonstrates the risks of human-mediated dispersion and selection. Invasive species are a major cause of crop loss and a global threat to food security (Blair and Hufbauer 2010; Paini et al. 2016). With increased geographic connectedness via trade, the threat of invasive species arriving to countries where they were previously absent is expected to increase (Saarman and Pogson 2015; Paini et al. 2016). In pest invasions, special attention has been given to newcomer species that can rapidly proliferate as they no longer face their natural enemies (50–52). However, given the evidence of bidirectional introgression following invasion, the importance of the local species should not be underestimated when developing effective biosecurity programs. Through introgression, local species can provide the invasive species with genes associated with local adaptation, thus contributing to its settlement. In addition, hybridization with the local species can reduce allee effects in the introduced species (7). Indeed, there is compelling evidence that species which hybridize have a higher likelihood of successful establishment (Currat et al. 2008; Saarman and Pogson 2015; Mesgaran et al. 2016).

In addition to human-mediated introduction of species, there is growing evidence that anthropogenically altered selection regimes drive exceptionally rapid adaptation (Song et al. 2011; Norris et al. 2015). Agricultural pests are subjected to massive insecticidal pressure, which drives selection for rapid development of resistance (Clarkson et al. 2014; Norris et al. 2015). Our observations suggest that increased insecticide spraying in Brazil to control *H. armigera* in the early stages of invasion acted as a selective force driving introgression of *CYP337B3*, by temporarily elevating the fitness of hybrids over that of the insecticide-susceptible *H. zea*. Furthermore, the selection regimes are rapidly changing in Brazil, and the widespread use of Bt-crops has considerably increased following the expansion of *H. armigera*. While there is field-evolved resistance to some *Bt* toxins for *H. zea* in North America, this appears to result from a down-regulation of midgut protease activity that reduces toxin activation and is associated with fitness costs (72, 73), so that, as noted above, it appears unlikely there will be any adaptive introgression of specific Bt-resistance-related genes into *H. armigera*, whose far greater genetic diversity already includes many specific resistance alleles with little or no fitness costs. In any case, the fate of the genes that cross species barriers will be also strongly affected by the demographic dynamics of these species which have substantially declined since 2012 along with other species of Noctuid moths (Santos et al. 2017; Piovesan et al. 2018; Fonseca-Medrano et al. 2019). These population declines seem to be driven by a large-scale selection regime linked to “El Niño-Southern Oscillation Effect” (ENSO) and other components of global change (Santos et al. 2017; Piovesan et al. 2018; Fonseca-Medrano et al. 2019). Thus, neglecting the interplay between selection due to local environmental heterogeneity in pesticide use, and selection imposed by different components of global change can obscure the prediction of the species’ evolutionary responses and the long-term dynamics of hybridization in the sympatric populations of these *Helicoverpa* moths.

Hybridization and introgression represent serious risks associated with invasive species and understanding the mechanisms that determine their impact is necessary for the effective management of biological invasions. Recent advances in whole-genome sequencing give us a chance to monitor the rapid evolution of invasive and local species with important practical implications to design insect resistance management (IRM) programs. As whole-genome data accumulate for biological invasions, the consequences of invasive species in adaptive evolution can be systematically characterized.

## Materials and methods

### Sample Collection and DNA Extraction

*Helicoverpa armigera* and *H. zea* moths were collected between 2012 and 2017 from three different localities in Brazil. The samples from 2012 and 2013 are referred to as “2013” and the samples collected between 2016-2017 were treated as “2017” or recent populations. The Brazilian samples included 67 *H. armigera*, 27 *H. zea* and 2 early hybrids closer to *H. zea,* from Bahia, Planaltina, and Mato Grosso states. As complementary groups for the admixture analyses, we included 4 *H. zea* pre-invasion from 2006 in Brazil, 8 *H. zea* from the United States, 9 *H. armigera armigera* from Asia, Europe, and Africa, and 14 *H. armigera conferta* and 6 *H. punctigera* from Australia which are all considered to be unrelated to the current *H. armigera* invasion in Brazil. Sample collection data are listed in Table S1-4. All samples were preserved in ethanol and RNAlater or stored at −20°C following collection. DNA was extracted using DNeasy blood and tissue kits (Qiagen) before quantification with a Qubit 2.0 fluorometer (Thermo Fisher Scientific).

### Whole genome resequencing and SNP Genotyping

Illumina libraries were produced following the manufacturer’s instructions, and 100-bp paired-end reads were generated (Illumina HiSeq. 2000, and Novogene, Biological Resources Facility, Australian National University, Australia). For BR-samples, libraries were prepared an adapted tn5 transposase protocol (Picelli et al 2014) and 150bp paired-end sequences were generated (Illumina HiSeq Xten, BGI, China) (Picelli et al. 2014: 5). Raw sequence reads were aligned to the *H. armigera* genome using BWA v 0.7.12. Reads were trimmed when quality in at least two bases fell below Q30 using VCFTOOLS v. 0.1.15, and only uniquely aligning reads were included in the analysis. Resulting BAM files were sorted before duplicate reads were annotated using Picard v. 1.138, and indexed with SAMTOOLS v. 1.3.1 (Li et al. 2009). HaplotypeCaller in GATK v. 3.7 (McKenna et al. 2010) was used to call SNPs in individual samples, and we used GenotypeGVCFs in GATK (McKenna et al. 2010) to estimate genotypes across all individuals simultaneously, implementing a heterozygosity value of 0.01. We used VCFTOOLS v. 0.1.15 (Danecek et al. 2011) to calculate mean coverage statistics for each sample. We further filtered our SNP dataset to include a minor allele count ≥2, mean site depth ≥5 and ≤200, missing data per site ≤0.50, and Phred-scale mapping quality ≥30; resulting in 17,257,450 genome-wide bi-allelic SNPs for the 150 individuals.

### Classification of parental and hybrids individuals

We conducted a genetic principal component analysis (PCA) and discriminant component analysis (DAPC) to visualize clustering of parental and hybrid genotypes using 50,000 SNP markers randomly sampled across all chromosomes with the packages “adegenet” (Jombart 2008), “Poppr” (Kamvar et al. 2014) and “vcfR” (Knaus and Grünwald 2017) in R v.3.3.2.

We further classified individuals as parental *H. zea* and *H. armigera,* hybrids or backcrosses based on the interspecific heterozygosity and the proportion of ancestry from parental species. We estimated the proportion of the genome that was ancestrally derived from *H. zea* using SNPs that strongly segregated between that species and *H. armigera armigera*. These diagnostic SNPs were determined via an association test in PLINK v1.90b (Purcell et al. 2007), taking only variants where p-value < 3.83 × 10^-14,^ thereby limiting the maximum number of alleles to fixed alleles from either *H. zea* or *H. armigera.* We used only pre-incursion allopatric populations of *H. zea* from Brazil and the USA and *H. armigera armigera* from the Old World for the association test resulting in ∼1,000,000 ancestry-informative SNPs. We calculated the proportion of ancestry and interspecific heterozygosity for the individuals using custom python scripts considering the sum of *H. zea* alleles divided by all alleles, and the proportion of heterozygous sites (https://github.com/wvalencia-montoya/helicoverpa-project).

### Genomic divergence between and within species

To estimate genome-wide differentiation we calculated *F_ST_* using the popgenWindows.py script from the repository (https://github.com/simonhmartin/genomics_general) in 100kb non-overlapping windows. We calculated *F_ST_* between allopatric populations of *H. zea* and *H. armigera* and sympatric post-incursion populations for the two temporal samples 2013 and 2017.

### Phylogenetic weighting

We used Twisst (Martin and Van Belleghem 2017) to explore evolutionary relationships across the genome of allopatric and sympatric recent populations of *H. zea* and *H. armigera*. We performed phasing and imputation of missing bases using Beagle (Browning and Browning 2007) with 10,000 bp step size and 100 kb overlapping sliding windows. Maximum likelihood trees for Twisst were generated with the phyml_sliding_window.py script (Martin and Van Belleghem 2017) with the GTR model in 50 SNPs sliding windows in PhyML (Guindon et al. 2010).

### Introgression across the genome

To test for the genome-wide extent of admixture resulting from hybridization between *H. armigera* and *H. zea*, we performed the D-statistic test implemented in AdmixTools v. 5.1 with a block jackknife size of 61 bp (Patterson et al. 2012). We considered that D-statistic estimates were indicative of significant introgression when z-scores ≥ 2.5, and thus with a p-value of ≤ 0.05. To estimate the genome-wide proportion of admixture we conducted an *F_4_* ratio incorporating *H. armigera conferta* from Australia as the fifth population (description of the model in Table S2) and using AdmixTools v. 5.1 with a block jackknife size of 61 bp (Patterson et al. 2012).

For *H. armigera*-like individuals we implemented a model phylogenetic model with the following branching: (((Old World *H. armigera*, Brazilian *H. armigera* Date 1 or Date 2) North American *H. zea) H. punctigera)* and (((Old World *H. armigera*, Australian *H. armigera conferta*) North American *H. zea) H. punctigera)* as a control (Fig. S1). For *H. zea* populations we considered the model (((North American *H. zea*, Brazilian *H. zea* Date 1 or Date 2) Old World *H. armigera), H. punctigera*). To detect introgression in specific regions across the genome we calculated *fd* in 100kb sliding windows with the ABBA-BABA.py scripts as in Martin *et al.* (Martin et al. 2015) and implementing the same topology models used for the D-statistic.

To further characterize introgressed regions from either parental species across the genome of individuals, we calculated the proportion of ancestry in sliding windows, using custom python scripts. The calculations incorporated the set of SNPs ancestry-informative SNPs with hybrid index averages across 100-kb non-overlapping windows. Sliding windows averages of the proportion of ancestry were plotted using custom R scripts, and using “ggplot2” (Wickham 2009), “Hmisc” (Jr and others 2019), and “Rmisc” (Hope 2013) packages.

To confirm introgression of *CYP337B3*, all individuals of *H. zea* 2017 were screened for the presence of this gene as in Joußen et al. (Joussen et al. 2012). Additionally, we mapped reads of recent individuals of *H. zea* to the complete coding sequence of *CYP337B3,* and confirmed presence with the Integrative Genome Viewer IGV_2.4.14 (Robinson et al. 2011).

### Tajima’s D

Tajima’s D was calculated in sliding windows of 10 kb in VCFTOOLS v. 0.1.15 (Danecek et al. 2011). We computed mean and standard deviation of Tajima’s D values for the Brazilian populations: *H. armigera* (2013), *H. armigera* (2017), *H. zea* (2013), *H. zea* (2017) including individuals with the *CYP337B3* haplotype and *H. zea* (2017), following (Norris et al. 2015). We plotted Tajima’s D using ggplot and ggalt along a region in chromosome 15, whereby *CYP337B3* gene extends from ∼11,436,000 to 11,440,000 bp.

## Acknowledgements

We are grateful to Simon H. Martin who provided key theoretical and practical advice on early stages of this study. Also, to Hilde Schneemman, Alejandra Duque, Federico Tamayo, and Bregje Wertheim who made insightful comments on the manuscript. We are also grateful to the cotton farmers in Western Bahia for their support during sampling collections, and ICMBio and MMA for the Authorizations for Scientific Activities (SISBIO no. 48218-3 and 38547/6). This study was funded by the Royal Society Exchange grant [IES\R3\170415] awarded to CJ and SE and the EMBO short-term fellowship [ASTF-6889] awarded to SE. TW, WTT, KHG and SE were supported by the CSIRO H&B Genes of Biosecurity Importance fund (R-8681-1). We are grateful to the Conselho Nacional de Desenvolvimento Científico e Tecnológico - CNPq for AS fellowship (process n°. 306601/2016-8) and research fund (process n°. 403376/2013-0), and Empresa Brasileira de Pesquisa Agropecuária - Embrapa (SEG MP2 n° 02.13.14.006.00.00). To Juanita Valencia for her support.

## Supporting information

**S1 Fig. Discriminant Component Analysis of *Helicoverpa* samples.** DAPC of *Helicoverpa armigera armigera (Haa), H. a. conferta (Hac),* and *H. zea (Hz)* populations resuming first 25 axes of variation.

**S2 Fig. Admixture results for *Helicoverpa armigera and H. zea.*** Admixture results for k=2, the number of clusters which receive more support.

**S3 Fig. Genome-wide *F_ST_* for different populations. A.** *F_ST_* for autosomes and Z-chromosome for different comparisons within and between species for allopatric and sympatric populations of *Helicoverpa armigera* and *H. zea.* **B**. *F_ST_* between *H. armigera* 2013 and *H. armigera* 2017.

**S4 Fig. Summary of the representation of the weights of the trees across the genome.** Proportion of the three possible unrooted relations between the four samples: ((Old World *H. armigera*, Brazilian *H. armigera* 2017) (New World *H. zea*, Brazilian *H. zea* 2017)). Blue represents the weight of the species topology ((Old World *H. armigera*, Brazilian *H. armigera* 2017) (New World *H. zea*, Brazilian *H. zea* 2017)). Green ((Brazilian *H. zea* 2017, Brazilian *H. armigera* 2017) (New World *H. zea*, Old World *H. armigera*)) and orange ((New World *H. zea*, Brazilian *H. armigera* 2017) (Brazilian *H. zea* 2017, Old World *H. armigera*)).

**S5 Fig. Tajimas’ D along the region in chromosome 15 containing *CYP337B3***. Tajima’s D along the region in chromosome 15 containing the *CYP337B* locus for populations of *H. zea* 2013 (blue) and *H. zea* in 2017 (green). Grey shading indicates the location of *CYP337B3* and the dashed line indicates Tajima’s D = 0.

**S1 Table.** *F_st_* within and between species for allopatric and sympatric populations of *Helicoverpa armigera* and *H. zea. H. armigera* (Old World) refers to the subspecies *H. armigera armigera* distributed in the Old World excluding Australia. *H. zea* (New World) comprises Brazilian pre-invasion and USA, and finally *H. armigera* 2013/2017 and *H. zea* 2013/2017 refer to the post-invasion sympatric populations in Brazil.

**S2 Table.** Results of F4-ratio tests which represents the mixing proportions of an admixture event. The populations are related to each other as portrayed in the model below. Alpha represents the proportion derived from *H. armigera armigera* from each subpopulation. Below, model explaining the F-4 ratio test. B’ and C’ are the populations that ae admixed in C. Adapted from Patterson et al. 2012 (Patterson et al. 2012).

**S3 Table.** Tajima’s D calculated across 10kb windows. Mean and standard deviation were calculated using all windows across the genome that had more than 100 SNPs. Tajima’s D at the window including the *CYP337B3* locus*. CYP337B3+* refers to all the *H. zea* individuals that contain the *CYP337B3+* gene.

**S4 Table.** Sample sequencing and collection data, proportion of hybrid ancestry, and presence of Sample sequencing and collection data, proportion of hybrid ancestry, and presence of *CYP337B3* for *Helicoverpa zea* 2017.

